# Individual-specific functional connectivity shows improved performance in detecting and predicting individualized symptoms of Alzheimer’s disease in elderly people with/without APOE ε4 allele

**DOI:** 10.1101/2022.11.14.516364

**Authors:** Lin Hua, Fei Gao, Xiaoluan Xia, Qiwei Guo, Yonghua Zhao, Zhen Yuan

**Affiliations:** Faculty of Health Sciences, E12 Building, University of Macau, Avenida da Universidade, Taipa, Macau SAR 999078, China; Centre for Cognitive and Brain Sciences, N21 Research Building, University of Macau, Avenida da Universidade, Taipa, Macau SAR 999078, China; State Key Laboratory of Quality Research in Chinese Medicine, Institute of Chinese Medical Sciences, N22 Research Building, University of Macau, Avenida da Universidade, Taipa, Macau SAR 999078, China

**Keywords:** Apolipoprotein E ε4 allele, individual-specific functional connectivity, atlas-based functional connectivity, Alzheimer’s disease, machine learning

## Abstract

To date, no reliable biomarkers are available that link individual-specific functional connectivity and patients’ individualized symptoms for early detection and prediction of Alzheimer’s disease (AD) in elderly people with specific genotypes. Meanwhile, functional magnetic resonance imaging (MRI) and machine learning are promising tools that can reveal the relationships between brain and behavior at individual level towards predicting the transition to AD. In this study, individual-specific functional connectivity was constructed in elderly participants with Apolipoprotein E (APOE) ε4 allele (*N* = 120) and without APOE ε4 allele (*N* = 115), respectively. In particular, machine learning based on a recursive feature selection technique was carried out to track multiple clinical symptoms among differing genotypes at individual level from normal aging (NA) and AD. It was found that the captured neuroimaging features in both APOE genotyping groups were able to distinguish the changes of clinical symptoms from NA to AD. Besides, our findings illustrated that the connections between individual-specific functional regions exhibited significantly higher correlation between estimated and observed scores in multiple clinical symptoms than those from atlas-based functional connectivity for both APOE genotyping groups, while no significant performance was detected when the data of two APOE genotyping groups were combined for the estimation models. Further, individual-specific between-network connectivity constitutes a major contributor for accessing cognitive symptoms in both APOE genotyping groups. Therefore, this study demonstrated the essential role of individual variation in cortical functional anatomy and the significance in combining brain and behavior for improving the accuracy in detection and prediction of AD in elderly people with specific genotypes.

## Introduction

Alzheimer’s disease (AD) is one of the most prevalent neurodegenerative disorders, causing the deficits in memory and executive function and difficulties in patients’ daily life (Haque & Levey, 2019). Accumulated evidence has demonstrated that AD progression is closely associated with specific genotypes (Karch & Goate, 2015; Lane-Donovan & Herz, 2017) and the spectrum of AD clinical phenotypes (Franzmeier et al., 2020; Mutlu et al., 2017). Meanwhile, it is challenging in carrying out early detection and prediction of AD at the individual level, particularly for patients with differing clinical phenotypes. Therefore, it is essential to inspect the brain-behavior relationship, identify the crucial biomarkers for the diagnosis of AD, and further develop effective prevention and treatment strategies for AD with specific genotypes.

Interestingly, human APOE genotype variants are one of the major genome-wide associated risk factors for AD in elderly people (Michaelson, 2014). The APOE gene has three gene alleles (APOE ε2, APOE ε3, and APOE ε4), thus producing six genotypes (ε2/ε2, ε2/ε3, ε2/ε4, ε3/ε3, ε3/ε4, and ε4/ε4). In particular, existing studies illustrated that the presence of ε4 allele dose-dependently enhances the risk of AD, with ε3/ε4 carriers and ε4/ε4 carriers showing 3.68-fold and 7-fold increased risks as compared to ε3 homozygotes (e.g., Bertram, McQueen, Mullin, Blacker, & Tanzi, 2007). In addition to the high risk for AD, APOE ε4 carriers are inclined to exhibit significantly poorer scores in cognitive functions, episodic memory, executive function, and perceptual speed as compared to APOE ε4 noncarriers (X. Li et al., 2021; Song et al., 2020). Further, the brain networks in both APOE genotyping populations have been inspected, indicating that both APOE ε4 carriers and APOE ε4 noncarriers manifest the brain disconnection and decreased functional connectivity mainly in the default mode network (DMN) (Mentink et al., 2021; Zheng et al., 2021).

Elderly participants might exhibit varying symptoms in AD individually since they might be in different stages of cognitive declination or at different levels of disease progression. Therefore, identifying the brain biomarkers with sufficient accuracy to track particular symptoms at the individual level would fundamentally benefit the way AD is assessed and managed in clinical practice. In particular, neuroimaging like structural and functional magnetic resonance imaging (MRI), has enabled us to make significant advances in identifying the potential biomarkers of AD. Likewise, brain network analysis has been performed to detect the macroscopic features that can distinguish well between various categories of dementia (J. Zhao, Du, Ding, Wang, & Men, 2020). However, existing evidence have not yet been integrated into a set of regional interactions which can effectively track each elderly individual’s present AD-related symptoms and assist in the evaluation of individuals with different AD genotypes.

In addition, biomarkers detected from the circuit anomalies in AD have been compromised by the inconsistent brain activation regions identified across individuals. Therefore, stable parcellations of cerebral cortex (Glasser et al., 2016; Gordon, Laumann, Gilmore, et al., 2017; Yeo et al., 2011) and subcortical structures (Ji et al., 2019; Wig et al., 2014) are essential for the large-scale investigation into human brain architecture, which provides a cortical taxonomy for inspecting regional or network-level alterations in cortical functions associated with AD. However, the downside of these atlases is that they only offer the functional organization of the brain at the population level rather than the peculiarities of a particular individual. To date, a majority of neuroimaging studies still relied on group-level atlases for accessing individual-level functional data (Mentink et al., 2021; J. Zhao et al., 2020). Although these atlas-level analyses were able to identify the relationships between brain connectivity and clinical demographic data (Langs et al., 2016; Mueller et al., 2013), some nuanced yet critical information was still missing since the symptoms associated with functional networks are highly variable across individuals. For example, recent studies showed that a number of essential characteristics of brain networks might be missing by the use of atlas-based templates (Braga & Buckner, 2017; Gordon, Laumann, Gilmore, et al., 2017). Consequently, these findings might dilute the brain-behavior correlations that are crucial for fully understanding of neural mechanism underlying AD when using the group-level atlas on individual participants.

In this study, a novel individual-specific strategy (D. Wang et al., 2015) that examined the individual differences in cortical functional architecture was proposed to improve the robustness and inter-individual reliability of functional connectivity analysis and to generate the relationship between the brain networks and participants’ individualized symptoms. Meanwhile, it is hypothesized that the use of individual-specific functional connectivity can detect effective biomarkers of clinical symptoms to facilitate early detection and predication of AD in both APOE ε4 groups. To test the hypothesis and quantify the relationship between brain and clinical symptoms, we examined the individual-specific functional connectivity of a large cohort of APOE ε4 carriers and noncarriers with varied degrees of cognitive symptoms, respectively. Specifically, machine learning was carried out to identify the connections that track multiple domains of cognitive symptoms [e.g., Mini-Mental State Examination (MMSE) and Immediate Recall Total Score (LIMM)] in different APOE ε4 groups. This study would demonstrate the critical contribution of accounting for individual variation in cortical functional anatomy to tracking multiple clinical symptoms and genotypes, which could open a new avenue for the diagnosis and prediction of AD.

## Materials and Methods

### Participants

Participants were retrieved from the phase 2 and phase 3 datasets from the Alzheimer’s Disease Neuroimaging Initiative (ADNI; https://adni.loni.usc.edu/) in light of the availability of T1 -weighted and resting-state functional MRI, APOE genotypes, and symptom severity assessment including MMSE and LIMM. According to APOE genotypes, participants were classified into two groups: 1) APOE ε4 carriers’ group with at least one APOE ε4 allele (genotype ε3/ε4 and ε4/ε4), 2) APOE ε4 noncarriers’ group with genotype ε3/ε3. Previous studies have verified that APOE ε2 allele could contaminate the dose effect of the APOE ε4 allele and result in different neuropathology (Berlau, Corrada, Head, & Kawas, 2009; Z. Li, Shue, Zhao, Shinohara, & Bu, 2020). Thus, individuals with ε2 allele (i.e., ε2/ε2, ε2/ε4, and ε2/ε3) were excluded due to the possible protective effect. In addition, neuroimaging data of participants with excessive head motions (see the third step of neuroimaging data preprocessing), severe artifacts, partial brain coverage, histories of obvious head trauma, and alcohol/drug abuse were also excluded for further analysis. Further, 235 elderly participants (120 APOE ε4 carriers and 115 APOE ε4 noncarriers) were selected for the present study (Table 1).

**Table 1.**
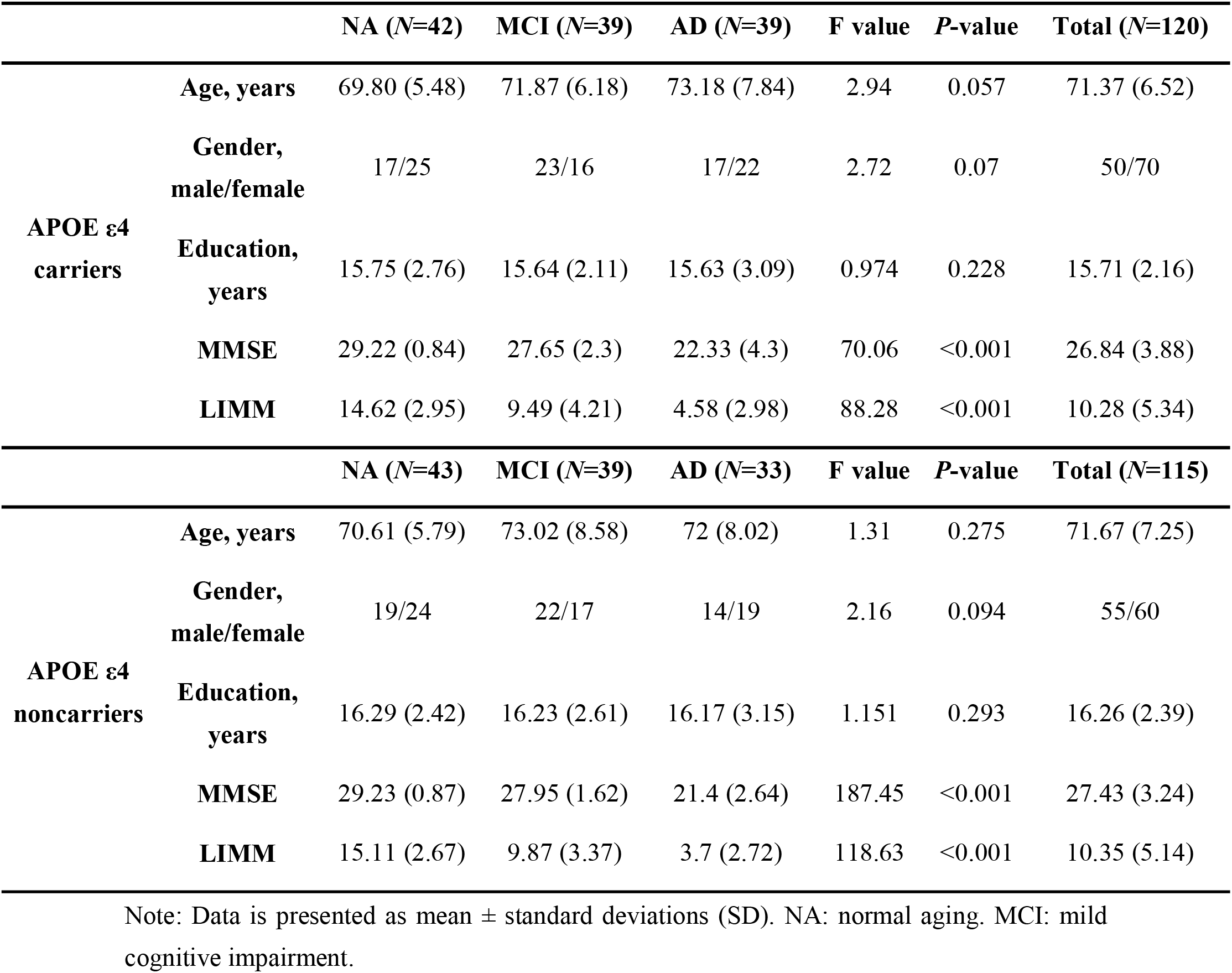
Demographic characteristics of all participants.

### APOE genotyping, neuropsychological assessment, and neuroimaging data acquisition

As previously described (Saykin et al., 2010), all participants’ APOE genotypes were screened by using DNA extracted from peripheral blood cells. The cells were collected in 10 ml EDTA plastic tubes and transported to the University of Pennsylvania AD Biofluid Bank Laboratory by overnight delivery at room temperature. More detailed information can be found in the ADNI Procedures Manual (http://adni.loni.usc.edu/methods/documents/).

A battery of neuropsychological assessments was completed by each participant in ADNI database. The present study focused on the results of MMSE and LIMM, which have been well validated and widely used as reliable tools for assessing cognitive impairment (Chapman et al., 2016). Importantly, the two assessments have been reported to be associated with brain functional connectivity (Edmonds et al., 2019; Ferreira et al., 2016). Table 1 presents the demographic and neuropsychological performances of all participants.

Both structural and functional MRI data of all participants were acquired on a 3-Tesla scanner from one of the following devices: General Electric (GE: Chalfont St. Giles, UK), Philips Medical Systems (PMS: Best, Netherlands), and Siemens (SIE: Erlangen, Germany). For structural T1-weighted images, GE scanners used inversion recovery-fast spoiled gradient recalled (IR-SPGR) sequences, while Philips and Siemens used magnetization-prepared rapid gradient echo (MP-RAGE) sequences. Related parameters were as follows: repetition time (TR) = 6.5, 7.4, and 2300 ms; echo time (TE) = 2.9, 3.1, and 3 ms; size = 1×1×1 mm^3^; flip angle = 9°, 11°, and 9°. Functional MRI (fMRI) data were collected using a gradient echo-planar pulse sequence with the following parameters: TR = 3000 ms; TE = 30 ms; flip angle = 90°; number of slices = 48; slice thickness = 3.4 mm. The first section (200 time points, 10 minutes) of fMRI data were extracted as each participant’s resting-state data. Detailed MRI scanner protocols for structural and functional sequences are available online (http://adni.loni.usc.edu/methods/documents/mri-protocols/).

### Neuroimaging data preprocessing

Resting-state fMRI (rs-fMRI) data were processed in terms of previous protocols (Buckner, Krienen, Castellanos, Diaz, & Yeo, 2011; D. Wang et al., 2020) that include the following steps: 1) removal of the first four frames; 2) slice timing correction with the FSL package (Smith et al., 2004) (http://www.fmrib.ox.ac.uk/fsl/); 3) rigid-body correction for head motion using FSL. Framewise displacement (FD) and root-mean-square of voxel-wise differentiated signal (DVARS) were then estimated using fsl_motion_outliers implemented in FSL. Volumes with FD > 0.2 mm or DVARS > 50 were marked as outliers (censored frames). One frame before and two frames after these outliner volumes were also flagged as censored frames, together with those lasting fewer than five contiguous volumes. Volumes with more than half labeled censored frames were removed; 4) linear regression of multiple nuisance regressors consisting of a vector of ones and linear trend, six motion correction parameters, averaged white matter signal, averaged ventricular signal, and temporal derivatives of the six motion correction parameters, averaged white matter signal, and averaged ventricular signal; 5) interpolation of censored frames with Lomb-Scargle periodogram; 6) band-pass filtering (0.009 - 0.08 Hz).

Structural MRI data were processed using the FreeSurfer 7.1.1 package (http://surfer.nmr.mgh.harvard.edu). The structural and functional images were aligned using boundary-based registration (Greve & Fischl, 2009). In a single interpolation, rs-fMRI data were aligned to a spherical coordinate system by sampling from the cortical ribbon. fMRI data of each individual was initially registered to the FreeSurfer surface template, which had 40962 vertices in each hemisphere. A 6 mm full-width half-maximum (FWHM) smoothing kernel was then applied to the fMRI data in the surface space. The smoothed data were then down-sampled to a mesh of 2562 vertices in each hemisphere using the mri_surf2surf function in FreeSurfer.

### Identifying functional ROIs in individuals

The functional regions of interest (ROIs) for individual participants were localized by using the previous methods (M. Li et al., 2019; D. Wang et al., 2015; D. Wang et al., 2020) with the following procedures:

Step 1. Individual participants’ cortical functional networks were mapped by using the iterative parcellation method (D. Wang et al., 2015). The algorithm was initially guided by the population-level functional network atlas constructed from 1,000 healthy participants. As the impact of the atlas on each individual brain parcellation was not identical for every participant or each brain area, the atlas was flexibly altered depending on the known distribution of inter-individual variability and the signal to noise ratio (SNR) distribution in a given participant. As the iteration progressed, the effect of population-based data diminished, enabling the final map to be totally driven by the data of each individual participant. More detailed information of the population-level functional network atlas and the iterative functional parcellation algorithm can be found in D. Wang et al. (2015).
Step 2. Using a clustering algorithm (mri_surfcluster in FreeSurfer software), the cortical networks of individual participants obtained from Step 1 were divided into a number of discrete patches. Each cortical network on the surface was spatially smoothed using a Gaussian kernel function (sigma = 1 mm) to reduce the influence of noise and matching costs. Only the template matching procedure (as explained below) would be impacted by the smoothing. The original unsmoothed area was kept for further analysis after a homologous ROI was detected.
Step 3. Individual participants’ discrete patches were matched to the 116 cortical ROIs generated from the population-level atlas that guided the search for a participant’s networks. The template matching procedure was performed for each cortical network as follows: 1) If an individual-level patch was overlapped (more than 20 vertices) with a single ROI in the population-level network, the patch was labeled as the same ROI in the atlas; 2) If a single individual-level patch was overlapped with numerous ROIs from a single network, the patch was divided into smaller patches. Vertices that were overlapped with the population-level ROIs were labeled together with these ROIs, producing the centers of numerous smaller patches. According to the geodesic distance on the brain surface, the remaining vertices in the original patch were allocated to the closest ROIs; 3) If a patch was not overlapped with any population-level ROI and the shortest distance between the patch and the ROI was less than a specific threshold, the patch was allocated to the ROI closest to it. The mean distance between any two vertices in the closest ROI was used as the specific threshold in the procedure; otherwise, the patch was labeled as “unrecognized”.

### Estimating within-network and between-network functional connectivity

Individual-specific functional connectivity was computed using Pearson correlation, resulting in a 116×116 connectivity matrix for each participant. For the comparison between individual-specific and atlas-based functional connectivity, Yeo’s group-level atlas (Yeo et al., 2011) was used to form a similar 116×116 connectivity matrix for each participant. Then, 116 ROIs based on both individual and atlas levels were divided into 18 networks (D. Wang et al., 2015). Functional connections were classified as within-network or between-network depending on whether they connected two ROIs in the same or different networks, respectively. Finally, each participant’s within-network and between-network connection values were calculated. The within-network connectivity was quantified by the averaged connectivity values of all ROI pairs within the same network, while the between-network connectivity was measured by the averaged connectivity values of all ROI pairs that involved one ROI within the specific network and the other ROIs outside this network.

### Estimating individual variability in size, position, and vertex-wise and ROI-based connectivity of the functional regions

To further examine the individual variability in the size and position of a functional region, as well as the vertex-wise and ROI-based connectivity were estimated using the similar strategy described in M. Li et al. (2019). The size of a functional region was estimated by the number of vertices that fell within this functional region. Then, the size variability for each ROI was quantified by the standard deviation of the size of functional regions across participants. The position of a functional region was determined by the coordinates of the center mass in this functional region. Then, the averaged geodesic distance between the ROI centers across participants was used to estimate the position variability for each ROI. Vertex-wise functional connectivity was denoted by the connectivity between vertices according to fsLR_32k surface mesh (59412 vertices). Furthermore, individual variability in ROI-based functional connectivity was computed as the strength of individual-specific ROI-to-ROI functional connectivity and atlas-based ROI-to-ROI functional connectivity across participants.

### Predicting individual symptom using individual-specific/atlas functional connectivity

Based on the connection between ROIs, a support vector machine for regression algorithm (L2-regularized L2-loss SVR model) from LIBLINEAR toolbox (https://www.csie.ntu.edu.tw-/~cjlin/liblinear/) was trained to predict each participant’s symptom severity ratings. The leave-one-out cross validation (LOOCV) method was utilized, in which data from N-1 participants were used to train the model. And then the model was applied to the data of the remaining participants to assess the severity of the participant’s symptoms. Before feature selection, covariates including age and gender were regressed from the features and the symptom severity ratings. The regressing weights were applied to the left-out dataset. To avoid over-fitting and remove duplicate information, features that exhibited strong associations with symptom ratings in the training dataset were chosen to train the model in each LOOCV (Finn et al., 2015). We used varied significance criteria in feature selection (*p* = 0.01, *p* = 0.005, and *p* = 0.001) to indicate big, medium, and small amounts of characteristics, respectively. To calculate the predicted symptom ratings, the de-confounded features from the testing data were fed into the trained model. To assess the symptom scores of all individuals, the procedure was repeated N times (*N* = 120 or *N* = 115). The estimated and observed symptom ratings were then compared to reveal the correlation. The greatest correlation between predicted and actual symptom ratings was given for various feature selection levels. The significance of the correlation was assessed using permutation testing (1,000 permutations), which randomly reshuffled the observed symptom among the participants. The *p* value was estimated by calculating the percentage of the correlation value of permutation data higher than the correlation value of real data. The same procedures were performed on three criteria for feature selection, which reported the highest correlation in each permutation test to accommodate the numerous comparisons associated with multiple thresholds. All *p* values were corrected for multiple comparisons by using Bonferroni method.

### Determining the contribution of each ROI and network in symptom estimation

The weight of a connection in the SVR model was used to quantify its contribution. In the present study, brain connections above the 90th percentile of absolute weight in each symptom estimation were considered as the most reliable connections. Specifically, the SVR model in the training data generated a weight coefficient for each feature of each LOOCV fold. The weights of each connection were then averaged over all LOOCV folds to determine their contribution to symptom estimation. If a connection was not chosen as a feature in one-fold, its contribution in this fold was set to zero. A specific ROI’s contribution was estimated by summing up the contributions of all connections affecting that ROI.

Between-network connections were separated from within-network connections to quantify the contribution of each functional network to symptom estimation (identical with the method of estimating within-network and between-network functional connectivity mentioned previously), which then grouped the predictive connections into the 7 canonical functional networks. The weight of each network in symptom estimation was calculated by adding the weights of the predicted connections of the involved network.

### Visualization

All imaging results were mapped onto the inflated PALS cortical surface and visualized by using CARET software (http://brainvis.wustl.edu/wiki/index.php/Caret:Download). Circos (http://circos.ca/) was used to construct the connectograms depicting connections that contributed to symptom estimation (e.g., Figure 4B). Connections that contributed to symptom estimation were further split into those positively or negatively linked with the symptoms, which were elaborated in Supplementary Figure S5.

## Results

### Inter-subject variability in size, position, and functional connectivity of individuals’ brain regions

In light of an iteratively individual-specific functional network parcellation approach, 18 cortical networks were mapped in each individual and then derived 116 discrete ROIs from these networks. Individuals showed significant inter-individual variability in these functional ROIs (see Supplementary Figure S1 for example). For each participant, functional connectivity was calculated among these ROIs in order to investigate the relationship between brain and behavior with specific genotypes (Figure 1).

**Figure 1.**
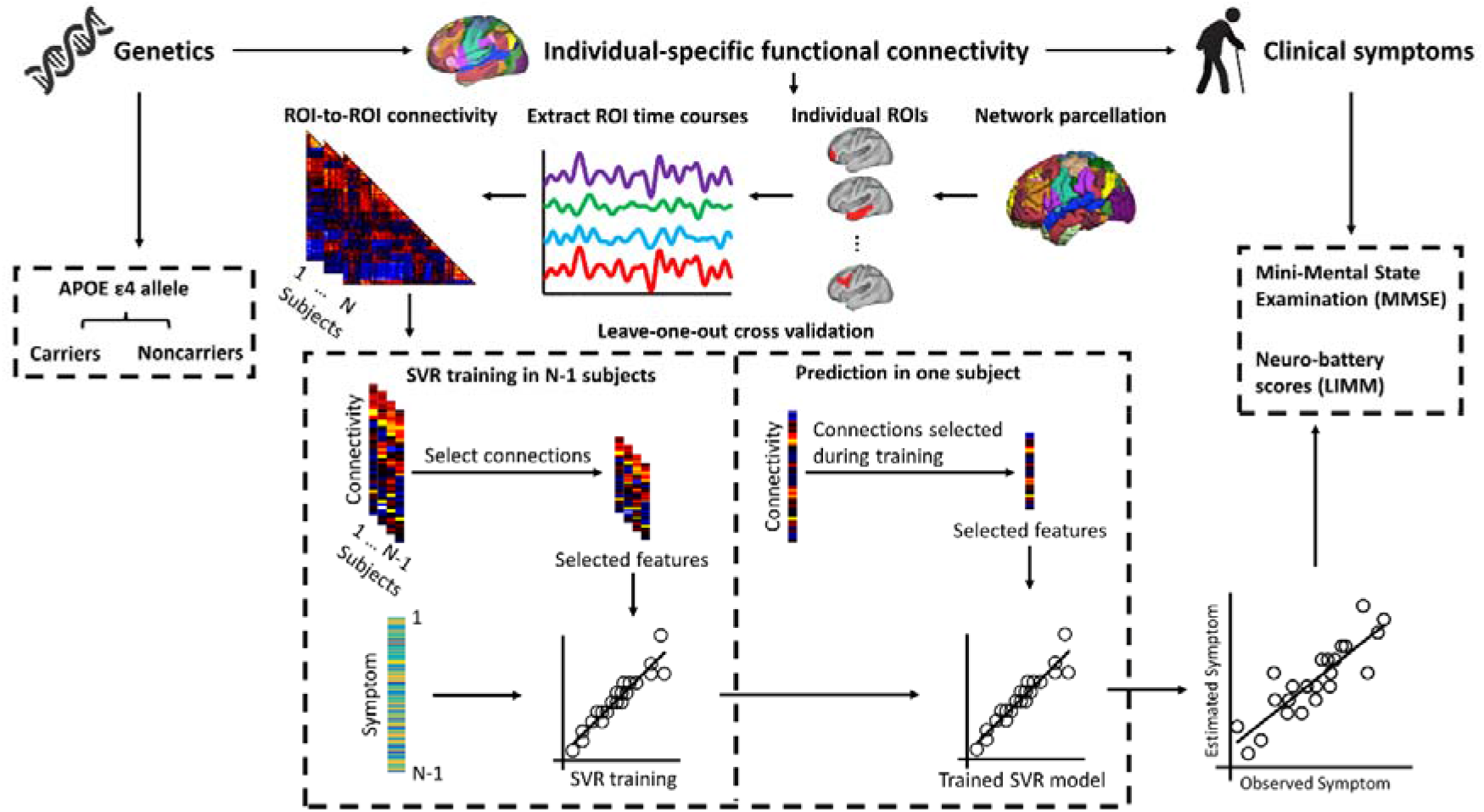
Procedure of estimating symptom scores in elderly people with/without APOE ε4 allele using functional connectivity among individually-specified ROIs. Participants were initially classified as APOE ε4 carriers and noncarriers depending on whether they carried at least one APOE ε4 allele, respectively. Then, based on the individual-level cortical network parcellation approach, we identified 116 homologous functional ROIs in each individual participant. The rs-fMRI signals in each ROI were then extracted and functional connectivity among these ROIs was computed, resulting in a 116×116 connectivity matrix for each participant. SVR model was trained to estimate each participants’ symptom scores based on ROI-to-ROI connectivity. To reduce the dimensionality of the input data, only a subset of connections that showed significant correlations with the symptom scores in the training dataset were selected as the relevant features to train the SVR. Data from *N*-1 participants were used to train the model and then the resulting model was applied to the data of the remaining participants to estimate the individual’s symptom. This procedure was repeated *N* times to predict the symptom scores of all participants. The correlation between the estimated and observed behavioral scores was then evaluated.

**Figure 2.**
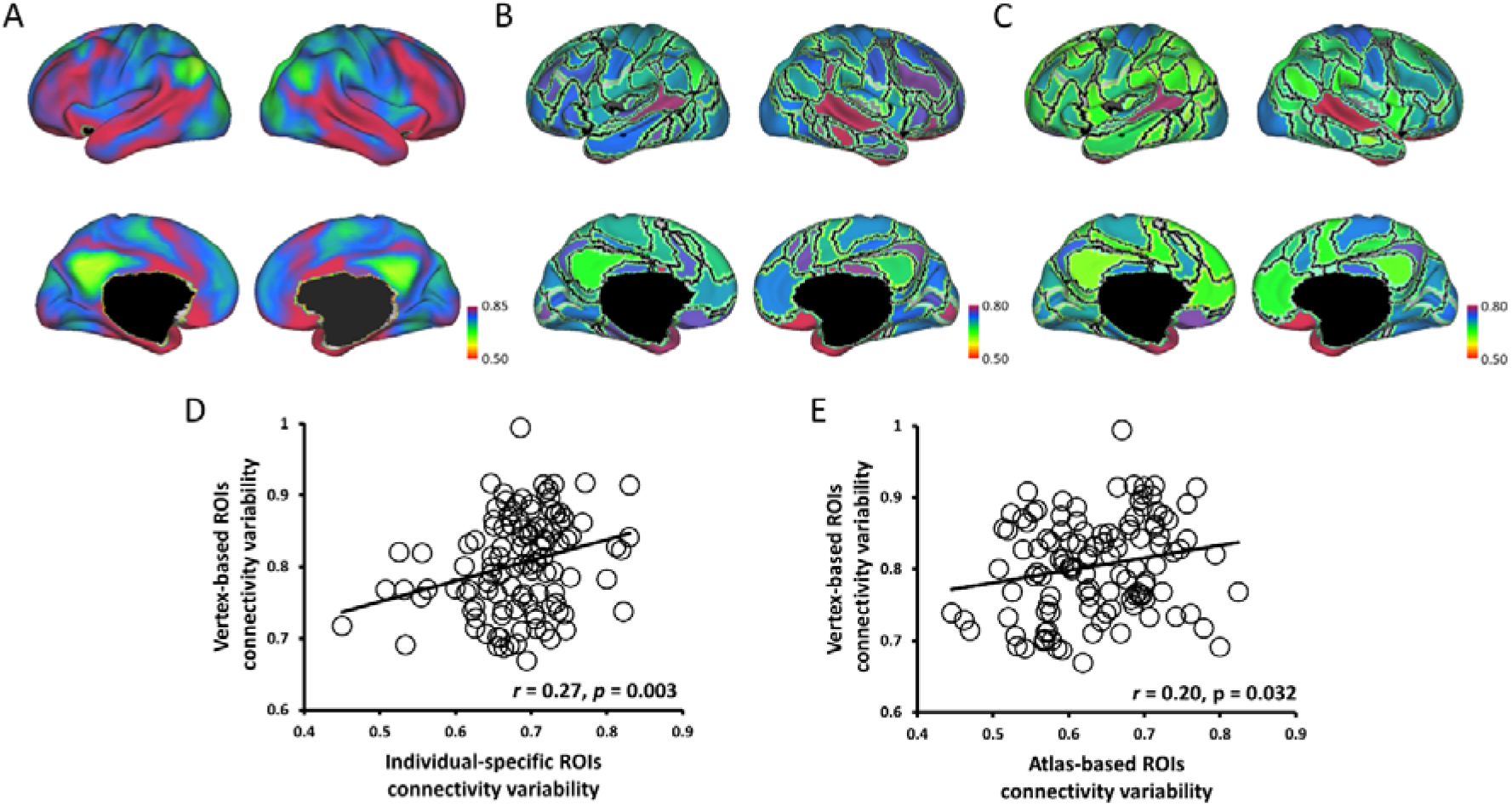
Individual-specific ROI connectivity was more highly correlated with vertex-based ROI connectivity than with atlas-based ROI connectivity. (A) Individual variability in resting state functional connectivity was calculated at each vertex across participants. Frontal and parietal cortices showed stronger individual variability than other cortices. (B) and (C) Individual-specific ROI connectivity and atlas-based ROI connectivity were quantified at 116 ROIs across participants. (D) and (E) Both individual variability in individual-specific ROI (*r* = 0.27, *p* = 0.003) and atlas-based ROI (*r* = 0.20, *p* = 0.032) connectivity showed a significant association with individual variability in vertex-based ROI connectivity.

**Figure 3.**
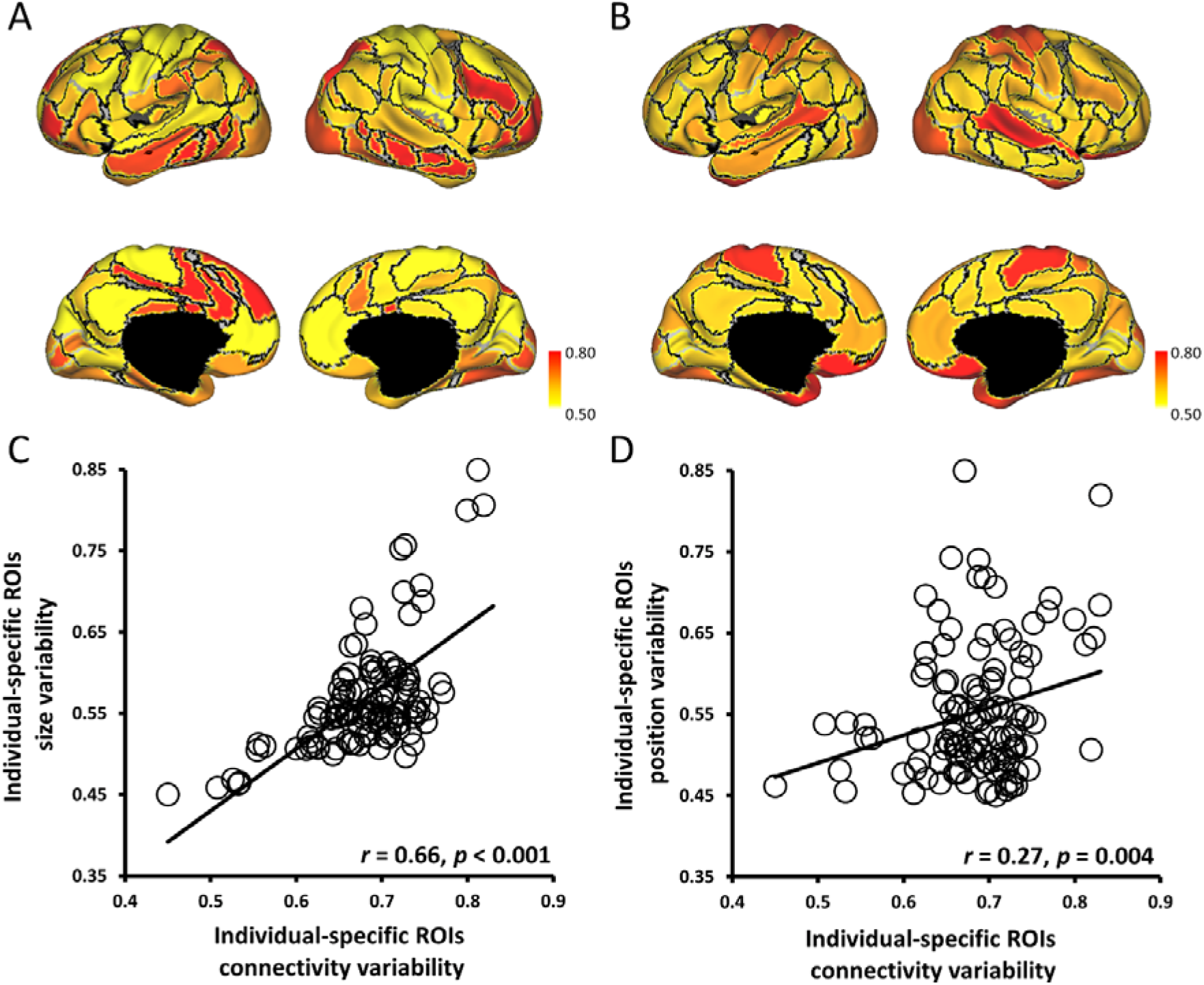
Individual variability in individual-specific ROI connectivity was significantly associated with individual variability in size and position of the functional regions. (A) and (C) Individual variability in ROI size was calculated for each of the 116 ROIs, and the size variability showed a significant correlation (*r* = 0.66, *p* < 0.001) with the variability in individual-specific ROI connectivity. (B) and (D) Individual variability in ROI position was also quantified for each of the 116 ROIs, and the position variability showed a moderate correlation (*r* = 0.27, *p* = 0.004) with the variability in individual-specific ROI connectivity.

To examine whether individual-specific approach carried high variation information across individuals, and individual-specific connectivity conducted a higher correlation to individual differences than atlas-based connectivity, this study quantified the individual variability in individual-specific connectivity, atlas-based connectivity, and vertex-based connectivity across all 235 participants. The relationship between individual-specific connectivity and individual variability in size and position of the 116 ROIs was then evaluated. Individual variability in vertex-based connectivity showed that functional connectivity was highly variable across individuals, especially in the frontal and parietal cortices which were associated with higher-order cognitive functions. Additionally, vertex-based functional connectivity was significant associated with the variability in connectivity strength among individual-specific ROIs (*r* = 0.27, *p* = 0.003) and atlas-based ROIs (*r* = 0.20, *p* = 0.032). Furthermore, individual-specific connectivity showed higher correlation with vertex-based connectivity than individual variability in atlas-based connectivity. Then, the individual variability in individual-specific connectivity was significantly associated with both variability in size (*r* = 0.66, *p* < 0.001) and variability in position (*r* = 0.27, *p* = 0.004). Therefore, these results could justify that individual-specific connectivity was more related to different aspects of individual variability than atlas-based connectivity, and the significant results in the current study corresponding to the individual-specific approach were caused by the individual variability across participants.

### Individual-specific functional connectome tracks MMSE symptoms

To determine whether individual-specific functional connectivity in different APOE genotyping carriers could track cognitive status, SVR model was trained to estimate the MMSE scores from each participant carrying or not carrying APOE ε4 allele. For the individuals with APOE ε4 allele, we found that a collection of functional connections among individual-specific ROIs were able to reliably predict MMSE symptom ratings. Meanwhile, the estimated and observed MMSE scores showed a modest yet statistically significant correlation (*r* = 0.41, *p* = 0.025, Figure 4A). Connections which exerted the greatest prediction on the MMSE symptom mainly employed the FPN and DMN (Figure 4B and S5A). In contrast, MMSE scores estimated by atlas-based functional connectivity identified in Yeo’s group-level atlas (Yeo et al., 2011) were not significantly correlated with the observed MMSE symptoms (*r* = 0.33, *p* = 0.087, Figure 4C). For the individuals without APOE ε4 allele, neither individual-specific ROIs (*r* = 0.23, *p* = 0.264) nor atlas-based ROIs (*r* = 0.21, *p* = 0.252) was able to estimate MMSE scores.

**Figure 4.**
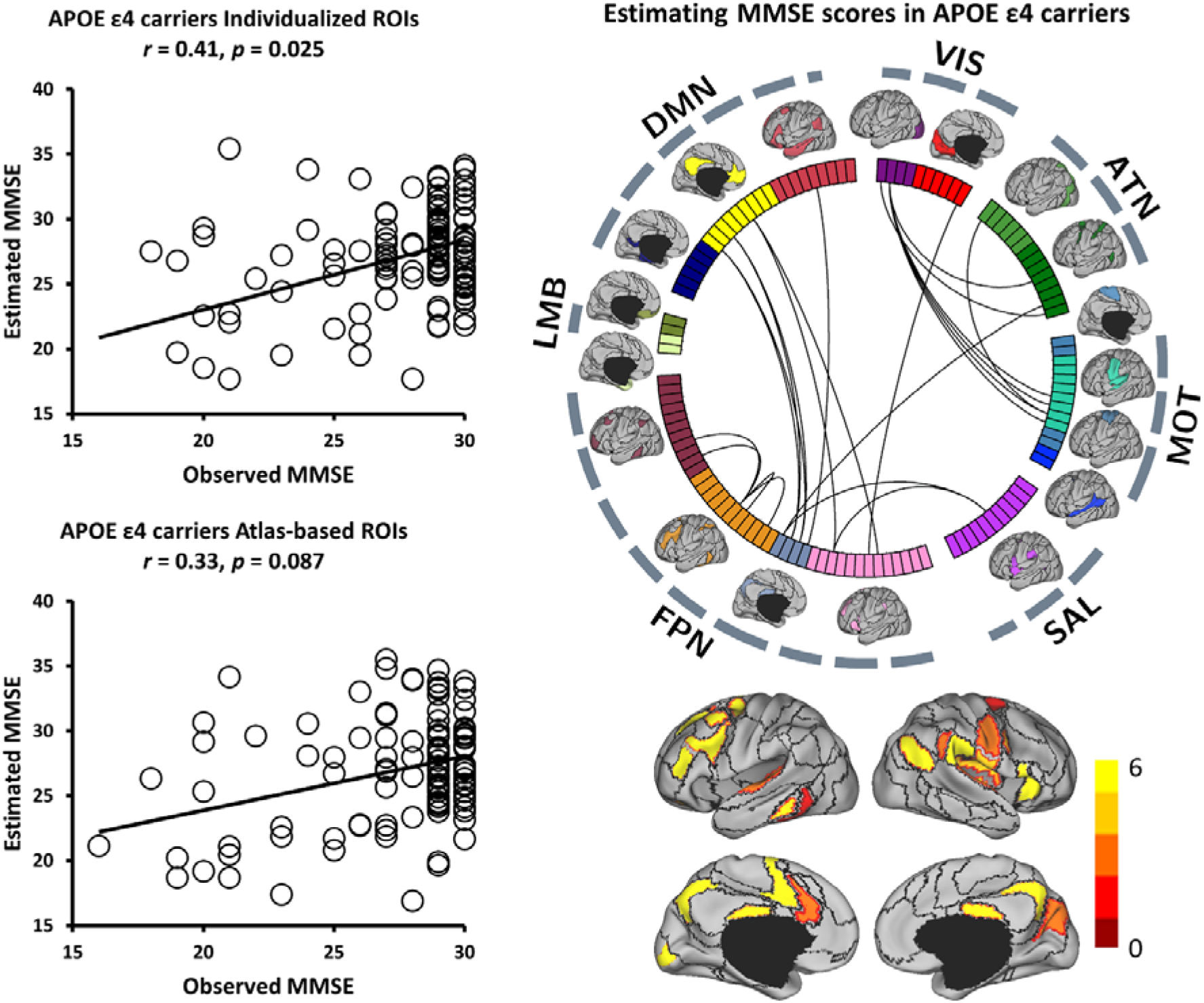
Functional connectivity among the individual-specific ROIs can better predict MMSE symptoms than that among the atlas-based ROIs in APOE ε4 carriers from normal aging to AD. (A) The scatterplot demonstrates the correlation (Pearson’s correlation, *r* = 0.41, *p* = 0.025) between the MMSE scores predicted by the connectivity among the individual-specific ROIs and the actually observed scores in APOE ε4 carriers from normal aging to AD. Each circle indicates a subject. Correlation significance was determined by using 1,000 permutations. (B) 116 ROIs derived from the 18 networks are presented by the colored rectangles under the corresponding brain networks. Twenty-two connections, which are above the 90th percentile of absolute weight in MMSE symptom estimation among APOE ε4 carriers, are specified by the black lines. Group-level maps of the 18 functional networks are shown on the cortical surface, respectively. ROIs involved in these predictive connections are plotted and color-coded with differing weights on the cortical surface below. (C) A similar analysis is performed using 116 ROIs defined in an atlas-based template. Functional connectivity among the atlas-based ROIs is not able to predict the MMSE scores in the group of APOE ε4 carriers (*r* = 0.33, *p* = 0.087). Correlation significance was determined by using 1,000 permutations.

Focusing on the symptom-related connections, we then examined whether the same connections defined by atlas-based ROIs would be less correlated with symptom scores. In other words, we tested if the connectivity features were already impaired by the atlas-based functional connectivity before the prediction model was applied. We found that the symptom-related connections (Figure 4B) were less correlated with symptom scores (Figure S3A) indeed, when defined by atlas-based ROIs. This finding suggested that the atlas-based ROIs obscured the symptom-related connections, thus further impeding symptom prediction.

### Individual-specific functional connectome tracks LIMM symptoms

To examine the specificity and precision of the current approach in estimating multiple symptoms, we next tested whether individual-specific functional connectivity can track LIMM symptom scores in the same group of individuals with or without APOE ε4 allele. For the group of APOE ε4 carriers, significantly positive correlation (*r* = 0.30, *p* = 0.048, Figure 5A) was again obtained between the estimated and observed LIMM symptom scores. Nevertheless, functional connectivity among the atlas-based ROIs cannot predict the LIMM scores (*r* = 0.23, *p* = 0.13, Figure 5C). Connections most contributing to LIMM symptom prediction mainly involved the FPN, DMN, and sensorimotor (MOT) (Figure 5B and S5B). For the APOE ε4 noncarrier group, functional connectivity among individual-specific ROIs was able to predict LIMM symptom scores (*r* = 0.44, *p* = 0.012, Figure 6A). Although atlas-based functional connectivity also showed a significant correlation between the estimated and observed LIMM scores, the correlation was relatively weaker as compared to the individual-based functional connectivity (*r* = 0.38, *p* = 0.041, Figure 6C). Connections most contributing to LIMM symptom prediction mainly involved the FPN, ATN, DMN, and MOT (Figure 6B and S5C). Additionally, decreased correlation with symptom scores were also found with the symptom-related connections (Figure 5B and 6B) defined by atlas-based functional connectivity (Figure S3B and S3C).

**Figure 5.**
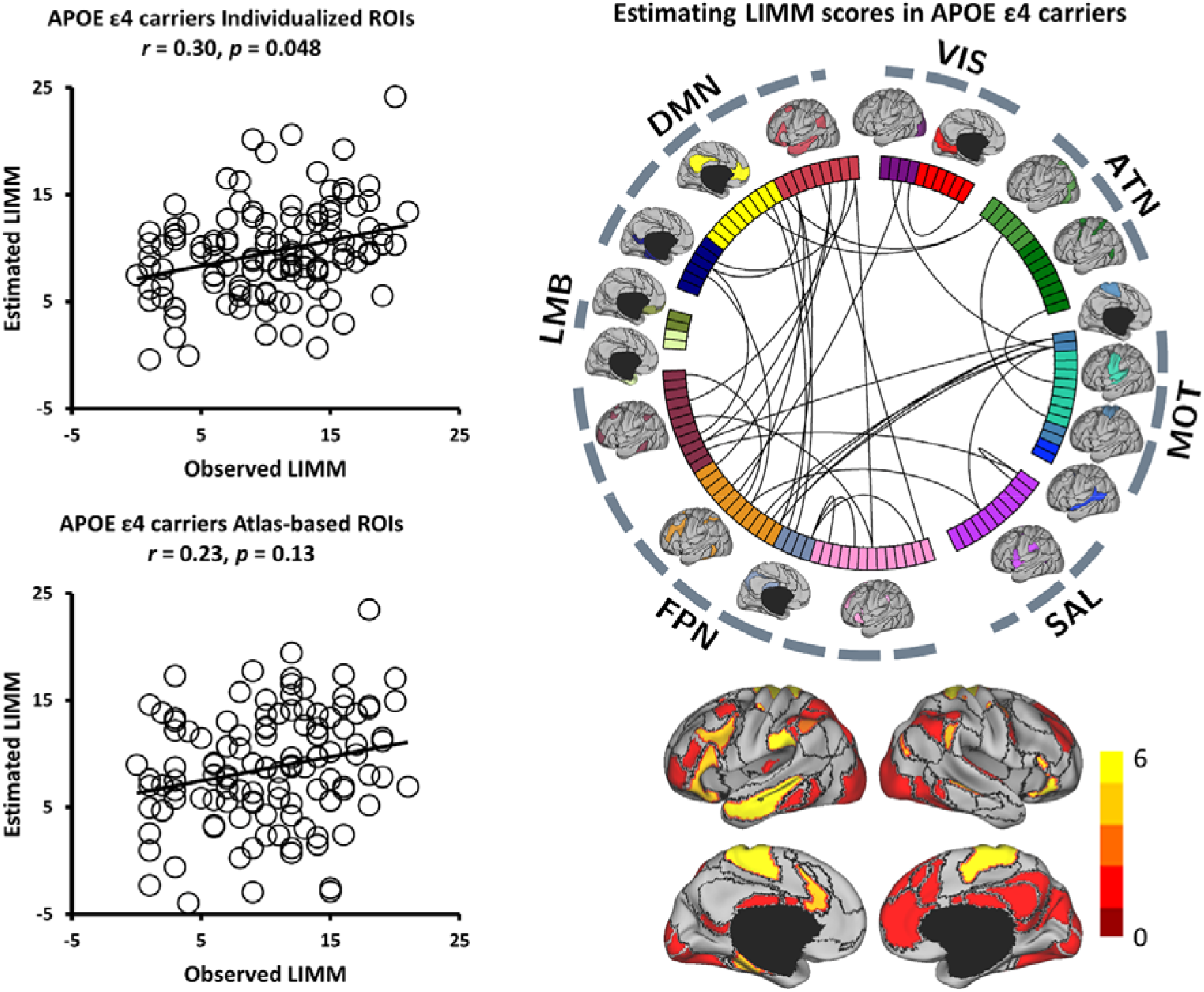
Functional connectivity among the individual-specific ROIs can better predict LIMM symptoms than that among the atlas-based ROIs in APOE ε4 carriers from normal aging to AD. (A) The scatterplot demonstrates the correlation (Pearson’s correlation, *r* = 0.30, *p* = 0.048, 1000 permutation test) between the LIMM scores predicted by the connectivity among the individual-specific ROIs and the actually observed scores. (B) Thirty-four connections, which are above the 90th percentile of absolute weight in LIMM symptom estimation among APOE ε4 carriers, are denoted by the black lines. ROIs involved in these predictive connections are plotted and color-coded with differing weights on the cortical surface below. (C) Functional connectivity among the atlas-based ROIs is not able to well predict the LIMM scores in the group of APOE ε4 carriers (*r* = 0.23, *p* = 0.13, 1,000 permutation test).

**Figure 6.**
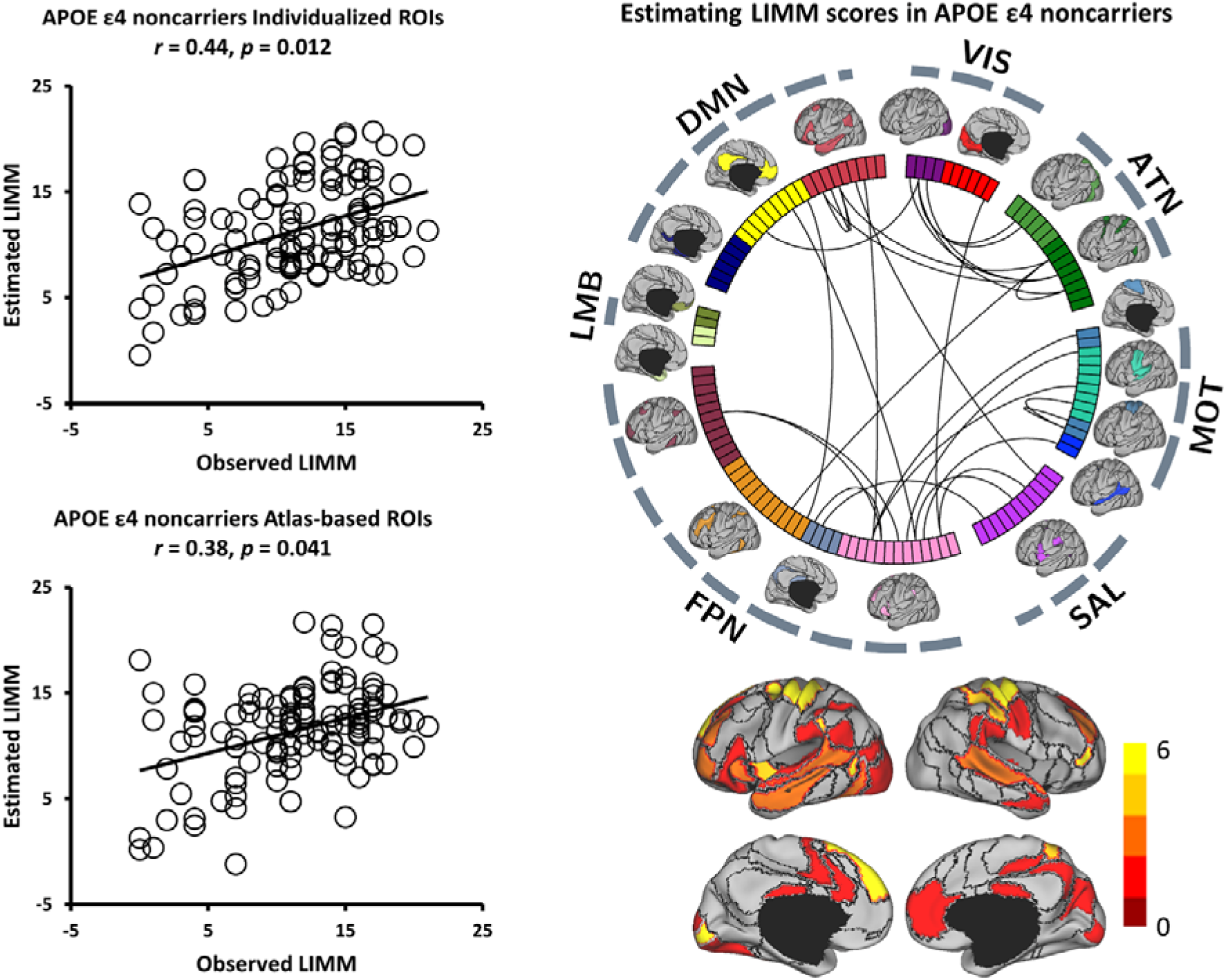
Functional connectivity among the individual-specific ROIs can better predict LIMM symptoms than that among the atlas-based ROIs in APOE ε4 noncarriers from normal aging to AD. (A) The scatterplot demonstrates the correlation (Pearson’s correlation, *r* = 0.44, *p* = 0.012, 1000 permutations) between the LIMM scores predicted by the connectivity among the individual-specific ROIs and the observed scores. (B) Twenty-nine connections, which are above the 90th percentile of absolute weight in LIMM symptom estimation among APOE ε4 noncarriers, are plotted by the black lines. ROIs involved in these predictive connections are plotted and color-coded with differing weights on the cortical surface below. (C) The correlation between the predicted and observed LIMM scores was weaker when connectivity was estimated using the atlas-based ROIs among these participants (*r* = 0.38, *p* = 0.041, 1,000 permutations).

### Between-network connectivity is a major contributor to the estimation of symptoms

By examining the functional connections that were predictive of MMSE and LIMM symptoms (e.g., Figure 4B, 5B, and 6B), we found that the majority of them were connections between functional networks rather than those within the same network. The strength of between-network connectivity showed a decrease of 7.25% on average when the ROIs were individual-specific compared to atlas-based (*p* < 0.01 for 17 of the 18 networks, paired t-test, Bonferroni corrected for 18 comparisons, Figure S2). Therefore, the reduced connectivity could result in more accurate symptom estimations, suggesting that between-network connectivity could be quantified more precisely when functional regions were localized in individuals.

Grouping the connections into the 7 canonical functional networks, the connections that contributed to MMSE symptom estimate model were mostly between-network connections including the FPN and DMN in APOE ε4 carriers (Figure 7). Furthermore, the connections that contributed to LIMM symptom estimation model were mostly between-network connections including the FPN, DMN, and MOT in both APOE ε4 carriers and APOE ε4 noncarriers (Figure 8). Specifically, ATN showed a higher contribution ranking in APOE ε4 noncarriers than in APOE ε4 carriers (Figure 8B).

**Figure 7.**
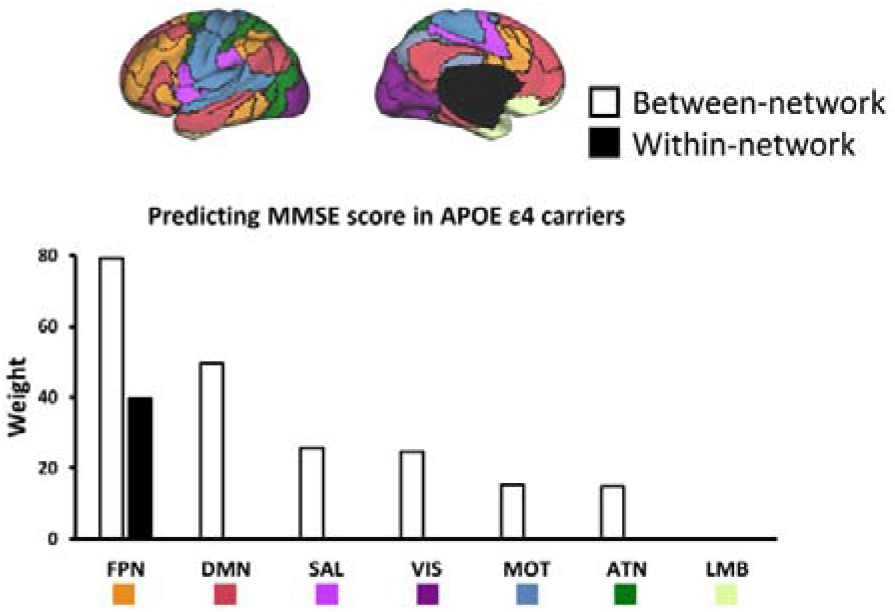
Between-network connectivity plays an essential role in predicting MMSE symptoms in APOE ε4 carriers. The functional connections most predictive of the MMSE scores in APOE ε4 carriers are grouped according to the 7 canonical networks. Connections contributing to the symptom prediction are mainly between-network connections (white bars). These between-network connections mainly involve the FPN and DMN.

**Figure 8.**
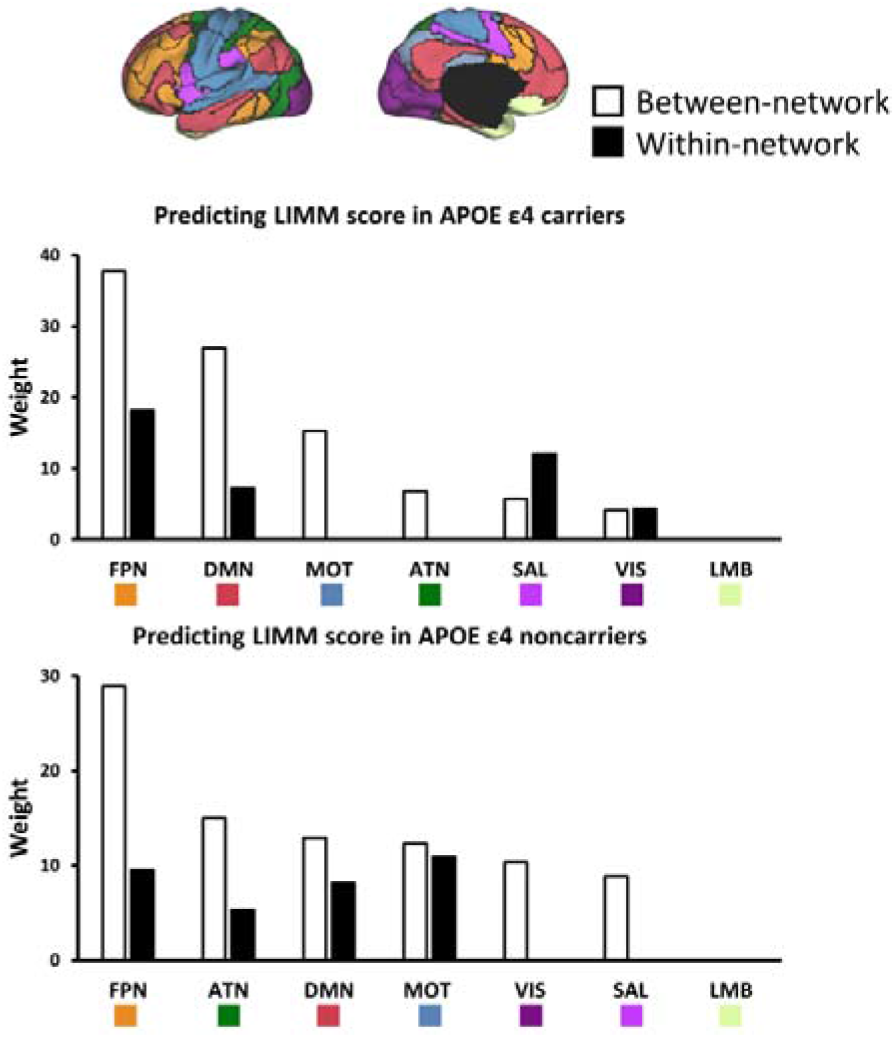
Between-network connectivity plays an essential role in predicting LIMM symptoms in APOE ε4 carriers and APOE ε4 noncarriers. (A) The prediction of LIMM scores in APOE ε4 carriers is mainly driven by between-network (white bars) involving the FPN, DMN, and MOT. (B) The prediction of LIMM scores in APOE ε4 noncarriers is mainly driven by the between-network involving the FPN, ATN, DMN, and MOT. Connections within the VIS and SAL also contributed to the prediction.

### Estimations of symptom dimensions perform better within different APOE genotyping groups

While a large body of imaging and neuropathology studies tended to suggest some distinct aspects of pathophysiology in elderly people with different APOE genotypes (Lane-Donovan & Herz, 2017; O’Donoghue, Murphy, Zamboni, Nobre, & Mackay, 2018), it is still unclear whether different functional connectivity could result in distinct neuropsychological representations in elderly people with differing APOE genotypes. To this end, we aggregated all 235 individuals across both APOE ε4 carriers and APOE ε4 noncarriers, and then trained SVR models to predict MMSE and LIMM symptoms on this merged dataset.

Although this aggregated analysis included the largest number of participants, individual-specific functional connectivity in this cross-genotype cohort cannot perform MMSE symptom estimation above the chance level (*r* = 0.18, *p* = 0.139, Figure S4A). Same results were also found in LIMM symptom estimation. Individual-specific functional connectivity failed to yield LIMM estimation which was not significantly correlated with the observed LIMM symptom rating (*r* = 0.23, *p* = 0.108, Figure S4B).

## Discussion

Drawing on the individual-specific functional network parcellation and machine learning approaches, the current study aimed at identifying the neuroimaging signatures in different APOE genotyping groups that could track cognitive symptoms from normal aging to AD. Our methods would enable data-driven estimations of the specific cortical connections which allows for the most accurate prediction of a given symptom score in elderly population with or without APOE ε4 allele. Specifically, the current results provide reliable readouts for testing how changes in a particular model affect the model’s ability to estimate a specific set of symptoms, which further establishes the relationship among brain functional connectivity and behavioral symptoms.

In line with previous studies (Braver, Cole, & Yarkoni, 2010; Gordon, Laumann, Adeyemo, & Petersen, 2017; Kanai, Bahrami, & Rees, 2010), individual variability results demonstrated that functional connectivity was highly variable across individuals, especially in the higher-order cognitive areas. The individual-specific approach used in current study can more precisely depict the individual differences than atlas-based approach. Furthermore, the comparison results indicated that compared to performing atlas-based functional connectivity, analysis carried out on individual-specific functional regions could promote the correspondence between functional connectivity and multiple symptom scores in differing APOE genotypes. This pattern was in line with the evidence from healthy population (M. Li et al., 2019) and psychotic illness patients (D. Wang et al., 2020), which showed that individual-specific functional connectivity manifested much greater accuracy to assess fluid intelligence and psychotic symptoms than using atlas-based template.

With regard to the connections most contributing to symptom prediction, previous studies reported that APOE gene-related symptoms could alter the resting-state functional connectivity among the frontal, temporal, and DMN regions (O’Donoghue et al., 2018; Z. Wang et al., 2019; Yu, Sporns, & Saykin, 2021). Yet, convergent evidence was unable to be obtained from normal aging to AD population (Jones et al., 2016; Koch et al., 2012), even though this is thought to be a gradual process from normal aging to mild cognitive impairment (MCI) and finally AD (Drachman, 2006). In the current work, we observed the most consistent results in APOE ε4 carriers that FPN and DMN connections contributed to MMSE symptom estimation model, while the FPN, DMN and MOT connections contributed to LIMM symptom estimation model in APOE ε4 carriers and noncarriers. Moreover, compared to the results in APOE ε4 carriers, the contribution ranking of ATN was higher in APOE ε4 noncarriers, thus suggesting that ATN might constitute a critical region corresponding to symptoms in APOE ε4 noncarriers. This finding is compatible with several proposals examining the abnormal circuit in MCI or AD patients relative to healthy participants (Z. Wang et al., 2019; C. Zhao et al., 2022).

Another noteworthy finding from the current research is that, in every symptom we tested, between-network connection in both APOE ε4 carriers and noncarriers implicated a critical role in predicting the severity of symptoms. Although the absolute values of between-network connectivity were significantly reduced after functional alignment, they still performed a more accurate prediction of symptoms. While exploring interactions across functional networks from normal aging to AD participants, previous studies primarily focused on the within-network variation as a key source of illness-related signal. In particular, the hyper- or hypoconnectivity within DMN was reported to be associated with the extent of cognitive decline (Jones et al., 2016). In this study, however, less significant relationship between within-network connectivity and clinical symptoms was identified, especially in the elderly participants with APOE ε4 allele. Since within-network variance may only reflect symptom-related general pathology of the disorder, the results of variation in between-network connectivity may represent the identification of symptom-related specific biomarkers. This finding is in line with recent studies suggesting that changes in between-network connectivity may signify the neuropathology changes in diseases (Satterthwaite et al., 2015; D. Wang et al., 2020). Importantly, our findings indicate that accounting for individual differences in functional network boundaries is critical, as mis-localization of networks might significantly obscure the actually low between-network correlations and further the identification of brain-behavior associations (Figure S4). To our knowledge, our study is among the first to combine individual-specific functional connectivity and machine learning approaches to examine the relationships among brain and behavioral symptoms in elderly people with specific genotypes from NA to AD. Therefore, the analysis framework of the current study can be extended to the investigations of brain-behavior associations in both healthy and clinical populations in the future.

Finally, some caveats need to be noted regarding the present study. Our study did not subdivide the groups of APOE ε4 carriers or noncarriers into normal aging, MCI, and AD since symptom rating might be low or invariant among individuals in the same subgroup. Meanwhile, subdividing groups might reduce the effective numbers of sampling and significantly bias the symptom estimation models. Moreover, we investigated the neuroimaging signatures of their correlation with symptom scores from APOE ε4 carriers and noncarriers, respectively, instead of differentiating the two groups. Finally, although feature selection was conducted using significant connectivity related to symptoms, the use of LOOCV still might increase the chance of overfitting. Future investigations could address these issues by subgrouping the participants in an increased sample size and performing various machine learning models to verify the estimated findings.

In summary, the current study found that the connectivity between individual-specific functional areas in elderly participants with different APOE genotypes was capable of yielding moderate-to-strong estimation levels across many primary categories of AD-related symptomatology. Notably, without accommodating individual differences in cortical functional architecture, conventional atlas-based functional connectivity was shown to be ineffective in predicting these symptoms in almost all cases. Furthermore, between-network variation explaining far more symptom-specific severity than in atlas-defined models, especially for the elderly people with APOE ε4 allele. Importantly, our research demonstrated the critical importance of accounting for individual variation in cortical functional anatomy in neurodegenerative research, which could be further extended to healthy population. Moreover, our work also highlighted the meaningful relationship among brain connectivity and symptoms, which can serve clinical utility in diagnosis, prognosis, and treatment for elderly people from potential to identified disorders.

## Acknowledgments

Data used in preparation of this article were obtained from the Alzheimer’s Disease Neuroimaging Initiative (ADNI) database (adni.loni.usc.edu). As such, the investigators within the ADNI contributed to the design and implementation of ADNI and/or provided data but did not participate in analysis or writing of this report. A complete listing of ADNI investigators can be found at: http://adni.loni.usc.edu/wp-content/uploads/how_to_apply/ADNI_Acknow-ledgement_List.pdf

## Supplementary material

Supplementary material is available in supplementary files.

## Data and Code Availability Statement

The data that support the findings of this study will be made available upon request to the corresponding author and the Alzheimer’s Disease Neuroimaging Initiative (ADNI) committee. The data from ADNI cannot be shared directly by the authors due to data protection defined by the ADNI data use policy (https://ida.loni.usc.edu/collaboration/access/appLicense.jsp).

## References

Berlau, D. J., Corrada, M. M., Head, E., & Kawas, C. H. (2009). APOE ε2 is associated with intact cognition but increased Alzheimer pathology in the oldest old. Neurology, 72(9), 829–834.

Bertram, L., McQueen, M. B., Mullin, K., Blacker, D., & Tanzi, R. E. (2007). Systematic meta-analyses of Alzheimer disease genetic association studies: the AlzGene database. Nature genetics, 39(1), 17–23.

Braga, R. M., & Buckner, R. L. (2017). Parallel interdigitated distributed networks within the individual estimated by intrinsic functional connectivity. Neuron, 95(2), 457–471. e455.

Braver, T. S., Cole, M. W., & Yarkoni, T. (2010). Vive les differences! Individual variation in neural mechanisms of executive control. Current opinion in neurobiology, 20(2), 242–250.

Buckner, R. L., Krienen, F. M., Castellanos, A., Diaz, J. C., & Yeo, B. T. (2011). The organization of the human cerebellum estimated by intrinsic functional connectivity. Journal of neurophysiology, 106(5), 2322–2345.

Chapman, K. R., Bing-Canar, H., Alosco, M. L., Steinberg, E. G., Martin, B., Chaisson, C., … Stern, R. A. (2016). Mini Mental State Examination and Logical Memory scores for entry into Alzheimer’s disease trials. Alzheimer’s research & therapy, 8(1), 1–11.

Drachman, D. A. (2006). Aging of the brain, entropy, and Alzheimer disease. Neurology, 67(8), 1340–1352.

Edmonds, E. C., McDonald, C. R., Marshall, A., Thomas, K. R., Eppig, J., Weigand, A. J., … Bondi, M. W. (2019). Early versus late MCI: Improved MCI staging using a neuropsychological approach. Alzheimer’s & dementia, 15(5), 699–708.

Ferreira, L. K., Regina, A. C. B., Kovacevic, N., Martin, M. d. G. M., Santos, P. P., Carneiro, C. d. G., … Busatto, G. F. (2016). Aging effects on whole-brain functional connectivity in adults free of cognitive and psychiatric disorders. Cerebral cortex, 26(9), 3851–3865.

Finn, E. S., Shen, X., Scheinost, D., Rosenberg, M. D., Huang, J., Chun, M. M., … Constable, R. T. (2015). Functional connectome fingerprinting: identifying individuals using patterns of brain connectivity. Nature neuroscience, 18(11), 1664–1671.

Franzmeier, N., Neitzel, J., Rubinski, A., Smith, R., Strandberg, O., Ossenkoppele, R., … Ewers, M. (2020). Functional brain architecture is associated with the rate of tau accumulation in Alzheimer’s disease. Nature communications, 11(1), 1–17.

Glasser, M. F., Coalson, T. S., Robinson, E. C., Hacker, C. D., Harwell, J., Yacoub, E., … Jenkinson, M. (2016). A multi-modal parcellation of human cerebral cortex. Nature, 536(7615), 171–178.

Gordon, E. M., Laumann, T. O., Adeyemo, B., & Petersen, S. E. (2017). Individual variability of the system-level organization of the human brain. Cerebral cortex, 27(1), 386–399.

Gordon, E. M., Laumann, T. O., Gilmore, A. W., Newbold, D. J., Greene, D. J., Berg, J. J., … Sun, H. (2017). Precision functional mapping of individual human brains. Neuron, 95(4), 791–807. e797.

Greve, D. N., & Fischl, B. (2009). Accurate and robust brain image alignment using boundary-based registration. NeuroImage, 48(1), 63–72.

Haque, R. U., & Levey, A. I. (2019). Alzheimer’s disease: A clinical perspective and future nonhuman primate research opportunities. Proceedings of the national Academy of Sciences, 116(52), 26224–26229.

Ji, J. L., Spronk, M., Kulkarni, K., Repovš, G., Anticevic, A., & Cole, M. W. (2019). Mapping the human brain’s cortical-subcortical functional network organization. NeuroImage, 185, 35–57.

Jones, D. T., Knopman, D. S., Gunter, J. L., Graff-Radford, J., Vemuri, P., Boeve, B. F., … Jack Jr, C. R. (2016). Cascading network failure across the Alzheimer’s disease spectrum. Brain, 139(2), 547–562.

Kanai, R., Bahrami, B., & Rees, G. (2010). Human parietal cortex structure predicts individual differences in perceptual rivalry. Current Biology, 20(18), 1626–1630.

Karch, C. M., & Goate, A. M. (2015). Alzheimer’s disease risk genes and mechanisms of disease pathogenesis. Biological psychiatry, 77(1), 43–51.

Koch, W., Teipel, S., Mueller, S., Benninghoff, J., Wagner, M., Bokde, A. L., … Meindl, T. (2012). Diagnostic power of default mode network resting state fMRI in the detection of Alzheimer’s disease. Neurobiology of aging, 33(3), 466–478.

Lane-Donovan, C., & Herz, J. (2017). ApoE, ApoE receptors, and the synapse in Alzheimer’s disease. Trends in Endocrinology & Metabolism, 28(4), 273–284.

Langs, G., Wang, D., Golland, P., Mueller, S., Pan, R., Sabuncu, M. R., … Liu, H. (2016). Identifying shared brain networks in individuals by decoupling functional and anatomical variability. Cerebral cortex, 26(10), 4004–4014.

Li, M., Wang, D., Ren, J., Langs, G., Stoecklein, S., Brennan, B. P., … Liu, H. (2019). Performing group-level functional image analyses based on homologous functional regions mapped in individuals. PLoS biology, 17(3), e2007032.

Li, X., Song, R., Qi, X., Xu, H., Yang, W., Kivipelto, M., … Xu, W. (2021). Influence of cognitive reserve on cognitive trajectories: role of brain pathologies. Neurology, 97(17), e1695–e1706.

Li, Z., Shue, F., Zhao, N., Shinohara, M., & Bu, G. (2020). APOE2: protective mechanism and therapeutic implications for Alzheimer’s disease. Molecular neurodegeneration, 15(1), 1–19.

Mentink, L. J., Guimarães, J. P., Faber, M., Sprooten, E., Rikkert, M. G. O., Haak, K. V., & Beckmann, C. F. (2021). Functional co-activation of the default mode network in APOE ε4-carriers: A replication study. NeuroImage, 240, 118304.

Michaelson, D. M. (2014). APOE ε4: The most prevalent yet understudied risk factor for Alzheimer’s disease. Alzheimer’s & dementia, 10(6), 861–868.

Mueller, S., Wang, D., Fox, M. D., Yeo, B. T., Sepulcre, J., Sabuncu, M. R., … Liu, H. (2013). Individual variability in functional connectivity architecture of the human brain. Neuron, 77(3), 586–595.

Mutlu, J., Landeau, B., Gaubert, M., de La Sayette, V., Desgranges, B., & Chételat, G. (2017). Distinct influence of specific versus global connectivity on the different Alzheimer’s disease biomarkers. Brain, 140(12), 3317–3328.

O’Donoghue, M. C., Murphy, S. E., Zamboni, G., Nobre, A. C., & Mackay, C. E. (2018). APOE genotype and cognition in healthy individuals at risk of Alzheimer’s disease: a review. Cortex, 104, 103–123.

Satterthwaite, T. D., Vandekar, S. N., Wolf, D. H., Bassett, D. S., Ruparel, K., Shehzad, Z., … Gennatas, E. D. (2015). Connectome-wide network analysis of youth with Psychosis-Spectrum symptoms. Molecular psychiatry, 20(12), 1508–1515.

Saykin, A. J., Shen, L., Foroud, T. M., Potkin, S. G., Swaminathan, S., Kim, S., … Craig, D. W. (2010). Alzheimer’s Disease Neuroimaging Initiative biomarkers as quantitative phenotypes: Genetics core aims, progress, and plans. Alzheimer’s & dementia, 6(3), 265–273.

Smith, S. M., Jenkinson, M., Woolrich, M. W., Beckmann, C. F., Behrens, T. E., Johansen-Berg, H., … Flitney, D. E. (2004). Advances in functional and structural MR image analysis and implementation as FSL. NeuroImage, 23, S208–S219.

Song, R., Xu, H., Dintica, C. S., Pan, K.-Y., Qi, X., Buchman, A. S., … Xu, W. (2020). Associations between cardiovascular risk, structural brain changes, and cognitive decline. Journal of the American College of Cardiology, 75(20), 2525–2534.

Wang, D., Buckner, R. L., Fox, M. D., Holt, D. J., Holmes, A. J., Stoecklein, S., … Li, K. (2015). Parcellating cortical functional networks in individuals. Nature neuroscience, 18(12), 1853–1860.

Wang, D., Li, M., Wang, M., Schoeppe, F., Ren, J., Chen, H., … Liu, H. (2020). Individual-specific functional connectivity markers track dimensional and categorical features of psychotic illness. Molecular psychiatry, 25(9), 2119–2129.

Wang, Z., Qiao, K., Chen, G., Sui, D., Dong, H.-M., Wang, Y.-S., … Han, Y. (2019). Functional connectivity changes across the spectrum of subjective cognitive decline, amnestic mild cognitive impairment and Alzheimer’s disease. Frontiers in neuroinformatics, 13, 26.

Wig, G. S., Laumann, T. O., Cohen, A. L., Power, J. D., Nelson, S. M., Glasser, M. F., … Petersen, S. E. (2014). Parcellating an individual subject’s cortical and subcortical brain structures using snowball sampling of resting-state correlations. Cerebral cortex, 24(8), 2036–2054.

Yeo, B. T., Krienen, F. M., Sepulcre, J., Sabuncu, M. R., Lashkari, D., Hollinshead, M., … Polimeni, J. R. (2011). The organization of the human cerebral cortex estimated by intrinsic functional connectivity. Journal of neurophysiology.

Yu, M., Sporns, O., & Saykin, A. J. (2021). The human connectome in Alzheimer disease—relationship to biomarkers and genetics. Nature Reviews Neurology, 17(9), 545–563.

Zhao, C., Huang, W.-J., Feng, F., Zhou, B., Yao, H.-X., Guo, Y.-E., … Zhang, X. (2022). Abnormal characterization of dynamic functional connectivity in Alzheimer’s disease. Neural Regeneration Research, 17(9), 2014.

Zhao, J., Du, Y.-H., Ding, X.-T., Wang, X.-H., & Men, G.-Z. (2020). Alteration of functional connectivity in patients with Alzheimer’s disease revealed by resting-state functional magnetic resonance imaging. Neural Regeneration Research, 15(2), 285.

Zheng, L. J., Lin, L., Schoepf, U. J., Varga-Szemes, A., Savage, R. H., Zhang, H., … Liu, Y. (2021). Different posterior hippocampus and default mode network modulation in young APOE ε4 carriers: a functional connectome-informed phenotype longitudinal study. Molecular Neurobiology, 58(6), 2757–2769.

