## Supplemental Files for "Individual-specific functional connectivity shows improved performance in detecting and predicting individualized symptoms of Alzheimer’s disease in elderly people with/without APOE ε4 allele"

**Supplementary Materials**

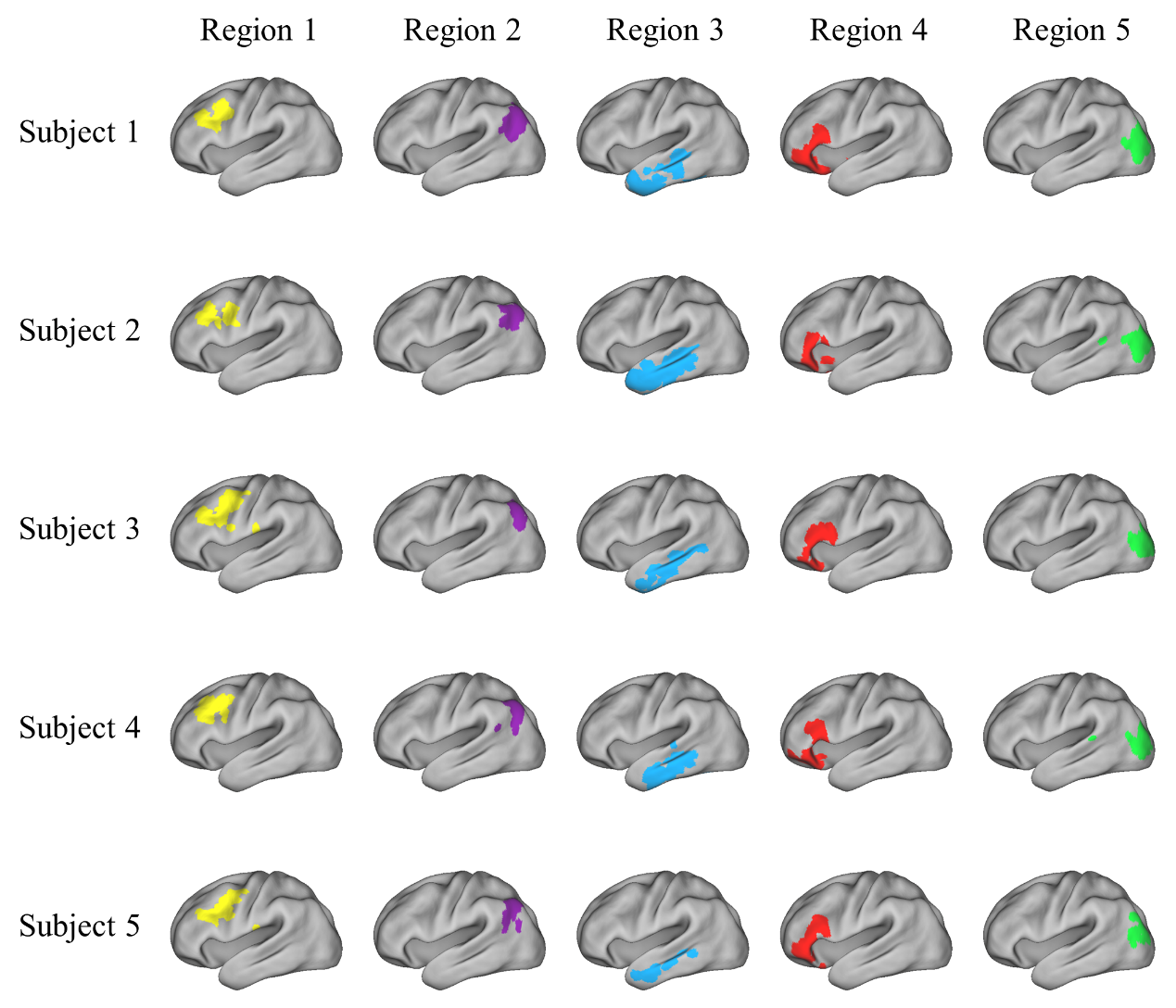

**Figure S1.** Five exemplary ROIs in five randomly selected participants were plotted on the brain surface. The ROIs showed marked variability in size and position. Colored regions denoted different ROIs across participants.

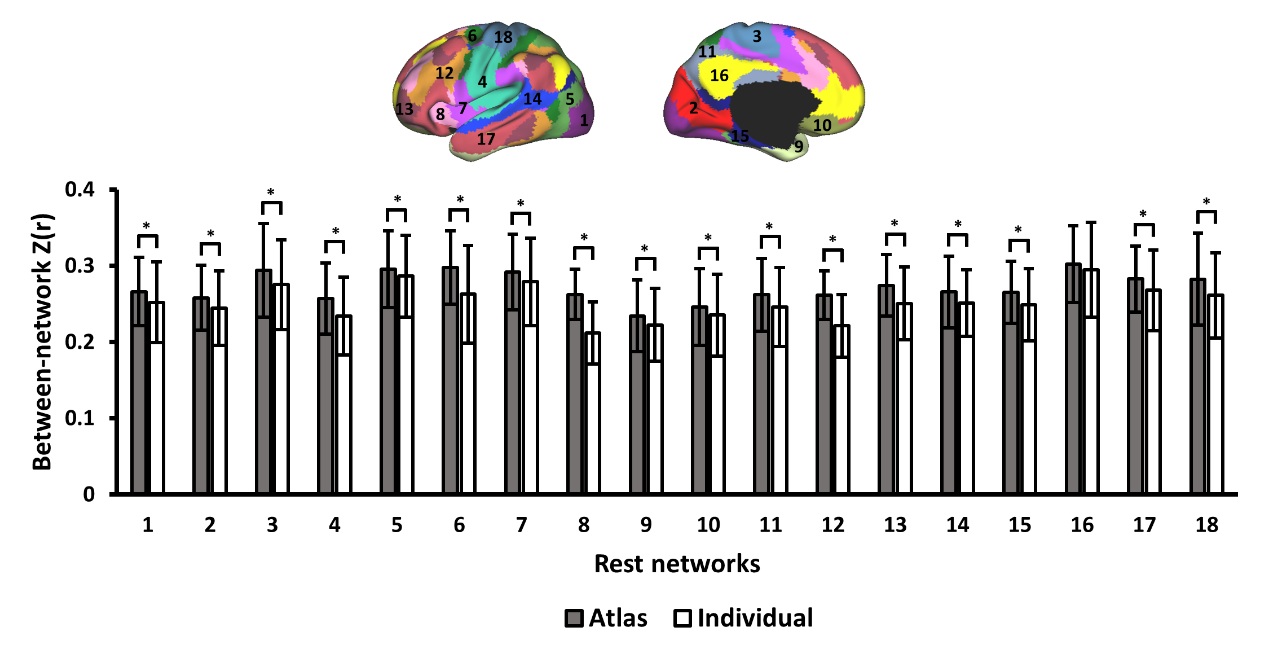

**Figure S2.** Functional connectivity strength between different networks was significantly reduced when the regions were individual-specific compared to atlas-based (*p* < 0.01 for 17 of the 18 networks. Error bars denotes SDs. * means *p* < 0.01, paired t-test, Bonferroni correction for 18 comparisons). Mean between-network connectivity decreased by 7.25% when the regions were individual-specific.

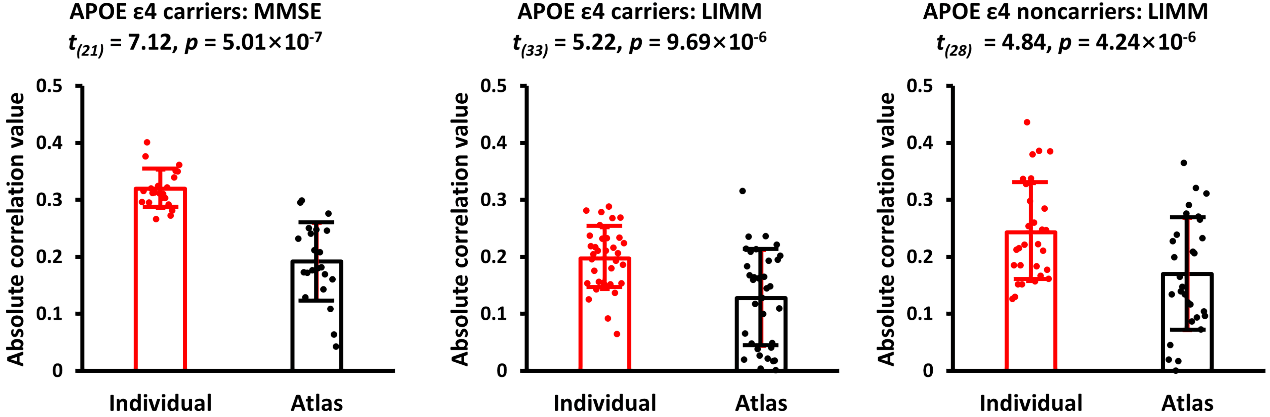

**Figure S3.** Individual-specific functional connections that showed a significant correlation with symptoms were redefined using the ROIs in the atlas. The same connections showed weaker correlations (paired t-test) with all symptom scores when the connections were defined using the atlas (black) compared to connections defined in individuals (red). All *p* values were corrected using Bonferroni method.

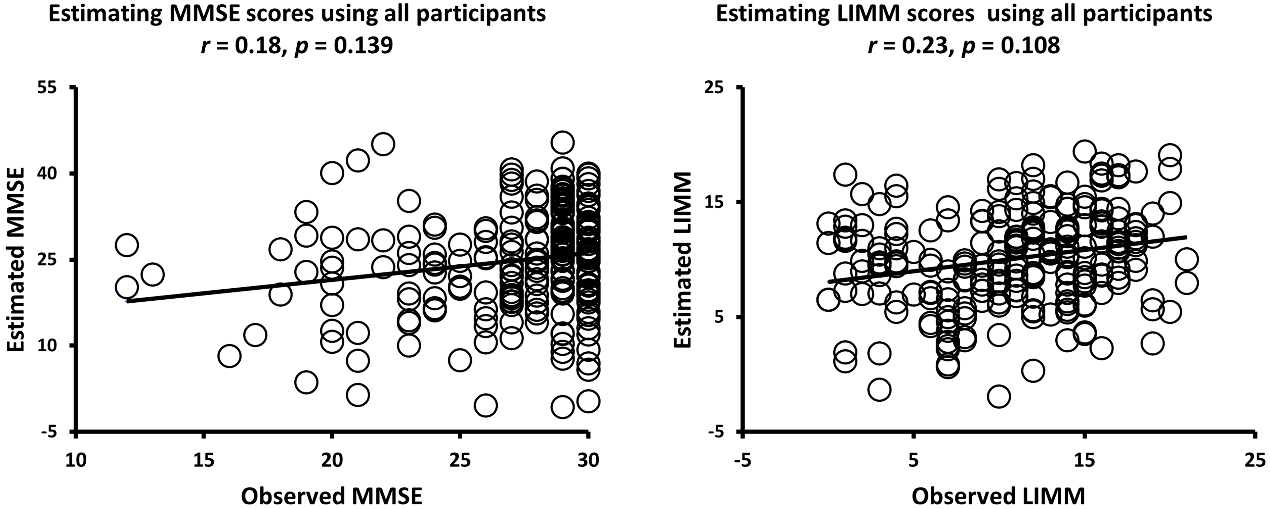

**Figure S4.** (A) Functional connectivity was not able to estimate the MMSE scores in the elderly participants across two APOE genotypes. (B) Functional connectivity was also not able to estimate the LIMM scores in the cross-genetic cohort.

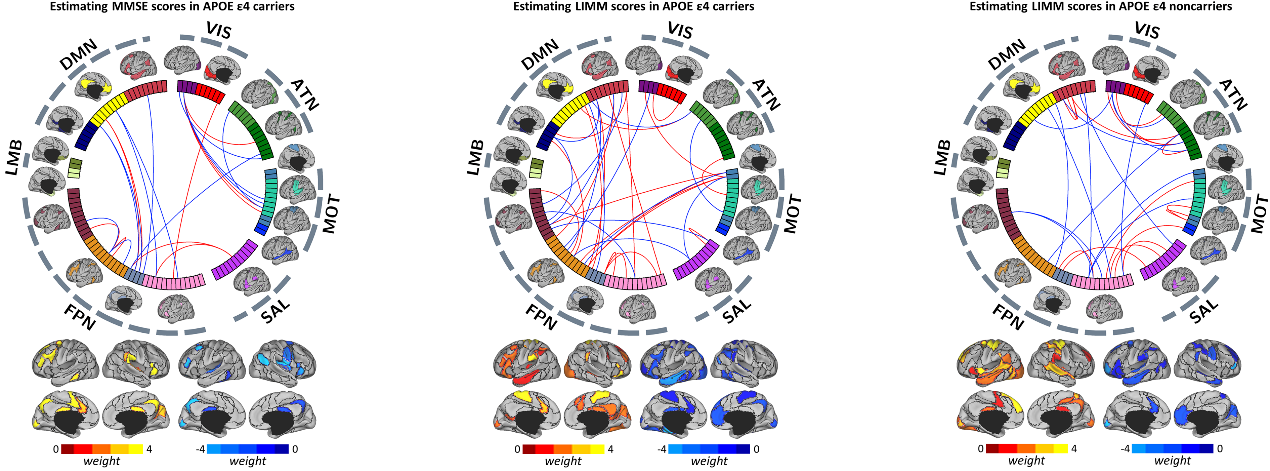

**Figure S5.** (A) Connections that contributed to the estimation of the MMSE scores in APOE ε4 carriers. (B) Connections that contributed to the estimation of the LIMM scores in APOE ε4 carriers. (C) Connections that contributed to the estimation of the LIMM scores in APOE ε4 noncarriers. Connections that were positively correlated with the symptom scores are shown in red and connections that were negatively correlated with the symptom scores are shown in blue.
